# Time-series RNA metabarcoding of the active *Populus tremuloides* root microbiome reveals hidden temporal dynamics and dormant core members

**DOI:** 10.1101/2025.02.19.639079

**Authors:** Jake Nash, Keaton Tremble, Christopher Schadt, Melissa A. Cregger, Corbin Bryan, Rytas Vilgalys

## Abstract

Dormancy is a key life history stage of many microbes that involves existence in a metabolically inactive state. Although dormant taxa may contribute little to community function, most microbial metabarcoding community surveys of microbial systems target DNA which captures dormant and dead taxa in addition to the active and living fraction of the community. RNA metabarcoding offers the potential to delineate the active fraction of microbial communities. Transitions between dormancy and activity may also serve as a rapid response mechanism for communities for communities to alter their function faster than turnover in community composition. Additionally, a focus on active microbiomes may provide further insight into which taxa should be prioritized as part of the core microbiome. We used DNA and RNA metabarcoding in samples collected as a time-series of the quaking aspen (*Populus tremuloides*) root microbiome across a natural environmental gradient to identify the degree of spatiotemporal dynamism in active and total microbial communities and to document the extent of dormancy in the core microbiome. We found that active bacterial and fungal communities were more temporally dynamic than total communities, while total communities exhibited a stronger response to spatial variability in site conditions. Additionally, we found that core microbiome members are frequently inactive, resulting in a reduced active core microbiome. This study stresses the need to focus on turnover in the active microbial community to detect variation in microbial communities in time-series and to use microbial activity levels as a key determinant of core microbiome membership.

## Introduction

Plant roots are host to assemblages of endophytic, epiphytic, and rhizospheric microbes that can facilitate root nutrient uptake, boost plant immunity, and increase resistance to abiotic stress [1, 2]. Large-scale plant root and soil microbiome studies have found that both soil chemistry and climate are strong controls on belowground microbial community composition [3–5]. In addition to the clear impact of soils and climate on microbial communities, individual extreme environmental events such as droughts, heat waves, and wildfires can cause rapid turnover in microbial community composition, likely due to variable stress tolerance by different community members [6, 7]. Microbes can also transition between states of metabolic dormancy or activity in response to environmental cues. The existence of metabolically inactive microbes has led some microbial ecologists to divide microbial communities into a *total community*, comprised of the full complement of microbes present in a sample, and an *active community*, comprised of the subset of microbes that are metabolically active [8, 9]. The ability of plant-symbiotic microbes to perform essential functions in plant root microbiomes such as nutrient uptake abiotic stress resistance depends on the ability of microbes to remain metabolically active under variable environmental conditions [10, 11]. Thus, it is essential to understand not only how environmental variation impacts *total communities*, but specifically how such variation affects the ecological dynamics of *active communities*. It has been shown that total microbial communities are structured by classic ecological processes such as ecological filtering, biotic interactions, and dispersal limitation [12, 13]. However, active community composition is likely mediated also by immediate environmental conditions that cause rapid transitions between dormancy and activity [8]. Yet, we lack an understanding of the importance of short-term environmental variation relative to longer term climatic variability in structuring active microbial communities. This knowledge gap prevents direct predictions of the influence of microbial communities on ecosystem processes which will be essential for the incorporation of microbial community data into ecosystem models.

Bacteria and fungi can rapidly transition between states of dormancy and activity dependent on suitable environmental conditions [14–16]. Dormant microbes are defined as cells exhibiting very low levels of metabolism and can either be resistant propagules such as spores and sclerotia, or vegetative cells with reduced metabolic activity and oftentimes accumulation of specific metabolites [8, 17]. Turnover of the composition of active microbial communities has the potential to occur more quickly than changes to the total community because it only requires spore germination or induction of metabolism rather than changes to population size through cell division or death [18]. Rapid changes in activity of bacterial and fungal symbionts can thus provide a means for microbial systems to quickly respond to acute environmental changes and the maintenance of essential metabolic processes.

In addition to the taxa in plant microbiomes that fluctuate in response to the environment, most microbial systems contain groups of taxa that are shared amongst distinct communities which are termed the “core microbiome” [19]. Microbes comprising the core microbiome are thought to be highly important in supporting host fitness and may be selectively recruited by their hosts [20]. The core microbiome is often quantitatively defined by microbes’ occupancy (the proportion of samples in which a microbe occurs), with occupancy thresholds often established of 0.95 or above to meet the criterion of “core.” Temporal stability in abundance is often also included as a criterion for core microbiome membership, thus requiring time-series sampling [19]. The ability of core microbial taxa to deliver essential functions to plants is likely dependent not only on their ability to persist across variable environmental conditions, but also to maintain metabolic activity under these environments and through time. However, few studies of core microbiomes have tested whether these taxa are consistently active through time and thus it is possible that certain “core” taxa are widely present as dormant propagules but have little functional importance for communities. Experimental assessments of the “active core microbiome” could thus further clarify which core microbiota are most important in supporting host acclimation to a wide variety of environmental conditions.

DNA metabarcoding is widely used to profile total microbial community composition through amplification of ribosomal marker genes. However, DNA metabarcoding can inflate the importance of dormant and dead microbes, which in some cases constitute up to 40% of the taxa detected with DNA metabarcoding [8]. In contrast to DNA-based studies, RNA metabarcoding is used to profile active microbial community composition and thus provide an indicator of potentially important taxa for ecosystem and host function. RNA metabarcoding studies rely on the amplification of cDNA generated from transcribed ribosomal genes, with the 16S subunit serving as a common target for bacteria and the ITS region serving as a common target for fungi [21, 22]. The ITS region is transcribed as part of a short lived primary rRNA precursor containing the small subunit, 5.8S region, and large subunit [23]. The transient nature of transcribed ITS regions means that it can serve as an indicator of cellular metabolic activity. However, because ribosomes themselves can persist in resting spores and other inactive cells, 16S RNA metabarcoding may have less power to discriminate the active from the total community [24, 25], which subtly distinguishes the insights available from ITS metabarcoding of fungi and 16S metabarcoding of bacteria.

Tree in the genus *Populus* are important model species for understanding plant responses to abiotic stress and plant-microbe interaction [26–28]. *Populus* spp. are dual-mycorrhizal, able to associate with both arbuscular mycorrhizal (AM) and ectomycorrhizal (ECM) fungi), as well as the melanized dark-septate endophytes (DSEs) that colonize roots without obvious symptoms [29, 30]. The surface (rhizoplane), interior (endosphere), and surrounding soil (rhizosphere) of roots is also colonized by a diverse bacterial community dominated by Proteobacteria, Actinobacteria, Bacteriodetes, and Firmicutes [29, 31, 32]. These bacteria express a wide array of molecular functions including production of antibiotics, secondary metabolites, plant hormones, and siderophores [33].

We utilized time-series sampling that leveraged the stark spatial and seasonal environmental variation in *P. tremuloides* stands in the Uinta Mountains in northeastern Utah to test the following hypotheses. We hypothesized that 1) active microbial communities will be more spatially and temporally dynamic than total microbial communities, 2) microbial taxa will exhibit unique responses to spatial and temporal environmental variation and 3) there will be a core microbiome comprised of highly metabolically active microbes across spatially variable sites that persists through time. In the Uinta Mountains, *P. tremuloides* occupies habitat types including mesic riparian sites, mid elevation semi-arid sites, and high elevation montane sites which we targeted in a hierarchical sampling design with stand-level ecological replication (see Fig. 1). Additionally, these sites are subjected to high levels of seasonal variation in moisture and temperature associated with predictable snow melt, drought, and monsoon periods that we leveraged with a time-series sampling design to incorporate temporal environmental variation.

**Figure 1.**
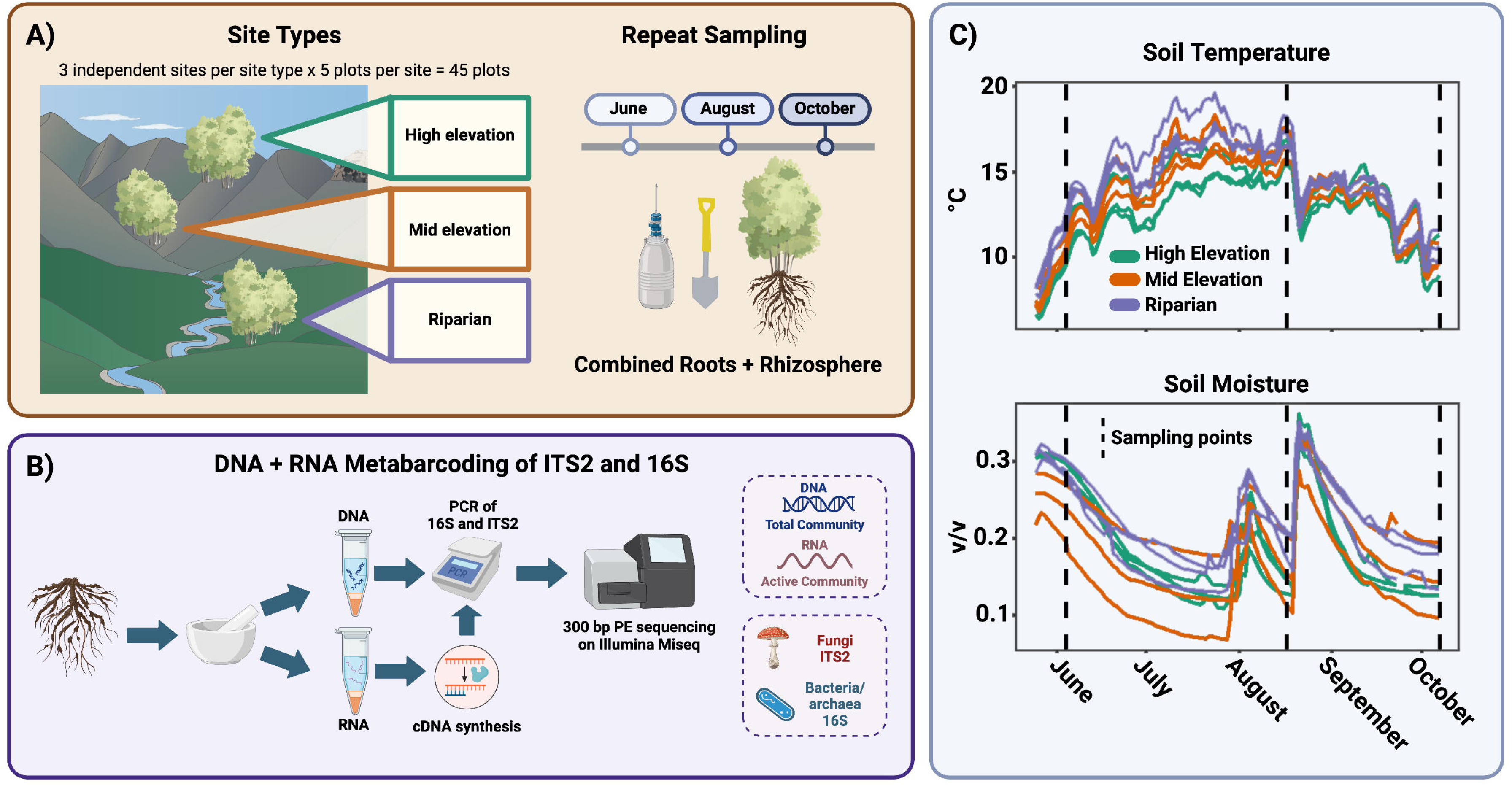
A) The field sampling design is depicted. A mixed root/rhizosphere sample was collected from three site types, with three independently replicated sites per site type. Each of these nine sites had five plots collected along a transect, for a total of 45 plots. These 45 plots were sampled three times in June, August, and October for a total of 135 samples. B) Samples were ground and subjected to parallel RNA/DNA metabarcoding for ITS2 (fungi) and 16S (bacteria/archaea) and sequenced on an Illumina Miseq with 300 bp paired-end sequencing. C) Average soil moisture and temperature for each site measured with in-ground sensors collecting data every 15 minutes.

## Methods

### Site description and sampling

We selected nine *P. tremuloides* stands located within a 6 km stretch along the Provo River valley in the Uinta–Wasatch–Cache National Forest across three habitat types– riparian, mid elevation, and high elevation (see Fig. 1). *P. tremuloides* constituted >90% of the tree stems at all sites to avoid confounding effects of co-existing overstory tree species. We confirmed the ecological distinctness of sites by collecting data on understory vegetation (see Fig. S1), soil chemistry (see Fig. S2), soil moisture/temperature sensors (see Fig. S3), and seasonal ion fluxes (see Fig. S4; Supplemental file S1). Riparian sites were located along 1-2 m wide streams feeding the Provo River, with sampling plots located within 2 m of the stream. Riparian sites had high understory cover of graminoids and (at one riparian site) *Equisetum* (see Fig. S1), elevated soil iron content, and the highest mean soil temperature (14.4°C; see Figs. 1, S3). Mid elevation and high elevation sites were both upland habitats located on northeast to northwest facing slopes. Mid elevation sites were intermixed with shrubs, whereas high elevation *P. tremuloides* sites were interspersed with tracts of coniferous (see Fig. S1). High-elevation sites had elevated cation exchange capacity (P < 0.05), slightly elevated soil organic carbon (n.s.; see Fig. S2), and the coldest soils (see Figs. 1, S3). Soil moisture was not significantly different between site types, but varied widely among individual sites (see Figs. 1, S3). Three independent, spatially separated sites were selected for each of the three site types. At each site, an 80 m transect was established, with five 2 m circular plots established along the transect with 20 m spacing that were selected to include a cluster of two to aspen stems. Aspen fine roots were sampled from each plot at three time points in a single growing season - early June, mid-August, and early October of 2021. Fine roots and adhering soil were sampled by digging within the top 15 cm of soil at the base of trees within the plot until approximately 2 g of root had been sampled. Samples were immediately flash frozen with liquid nitrogen in the field and kept at -80°C until processing.

### Molecular analyses and bioinformatics

Detailed molecular and bioinformatics are provided in Supplemental file S1, but a brief summary is provided here. DNA and RNA were extracted from each fine root sample, including adhering rhizosphere soil. RNA was reverse transcribed to generate cDNA which was then used in a PCR metabarcoding workflow along with the DNA to generate 16S (for bacteria and archaea) and ITS2 (for fungi) libraries with standard primers [29, 34, 35]. Although the 16S primers used target both bacteria and archaea, we refer to that dataset as the bacterial dataset in this manuscript because archaea were in very low abundance. Libraries were sequenced on two separate runs of an Illumina MiSeq with 300 base pair paired end sequencing with v3 chemistry at Duke University’s Center for Genomic and Computational Biology. Sequence data were processed with QIIME 2 v.2021.11 [36]and DADA2 [37] and assigned taxonomy with the UNITE v.8.3 99% database for fungi [38] and the Greengenes 13_8 99% database for prokaryotes [39].

### Statistics

16S and ITS data were rarefied to 13,407 and 17,182 sequences, respectively, to achieve an even sequencing depth while retaining as many samples as possible. Alpha diversity metrics were calculated using the *phyloseq* package [40]. The *lme4* package [41] was used to test for effects of site type, season, and metabarcoding method (RNA vs DNA) on alpha diversity with a random effect of site. Dissimilarity matrices were generated by performing square root transformation followed by Wisconsin double standardization before calculating Bray-Curtis dissimilarity and then used to test for the effects of site type, site, plot, season, and metabarcoding method on fungal and bacterial community composition using the adonis2() function in the *vegan* package [42] with 999 permutations. Beta-diversity patterns were visualized using NMDS conducted on the same dissimilarity matrices using the metaMDS() function with 100 iterations [42]. The envfit() function from *vegan* was used to test for effects of continuous environmental variables on microbial community composition [42]. The betadisp() function in *vegan* [42] was used to test for differences in beta dispersion across site types and season. LinDA [43] was used with a mixed model formula with site included as a random factor to identify differences in taxon abundance in response to dataset (RNA vs. DNA), site type, and season.

## Results

### Response of total and active communities to spatiotemporal environmental variation

DNA-based analyses showed a greater sensitivity to detect spatial variation in fungal and bacterial community composition due to site type, while RNA-based analyses better detected temporal variation due to season. Although community composition was more strongly structured by site type than by season for both fungal and bacterial communities (PERMANOVA; Tables S4, S5), this effect was stronger for DNA than RNA datasets. Conversely, seasonality had a greater effect on active community composition than on total community composition (PERMANOVA; see tables S6, S7, S8, S9). To confirm this pattern we identified the number of taxa which significantly varied in abundance by site or season. A greater number of taxa varied in abundance by site type versus season in the DNA dataset (20 vs. 55 fungal OTUs; 38 vs. 92 bacterial ASVs), while a greater number of taxa varied by season in the RNA dataset (40 vs. 7 fungal OTUs; 60 vs. 17 bacterial ASVs). There was little overlap in the assemblages of taxa found to be variable by site type or season between the DNA and RNA datasets (see Fig. 2). Most of the taxa that varied in abundance by site type reached peak abundance in the high-elevation sites in both the RNA and DNA datasets, indicating a larger number of high-elevation specialists. Amongst the high-elevation specialists (for which both DNA-based and RNA-based tests were significant), there were three ECM fungi (*Hebeloma crustulinforme*, *Hebeloma mesophaeum*, and *Geopora tolucana*), three potential DSEs in the Helotiales (*Cistella* sp., *Varicosporium* sp., and *Tetracladium* sp.), and three bacteria (Actinosynnemataceae sp., Cytophagaceae sp., and Chitinophagaceae sp.). Dominant fungal and bacterial lineages generally showed little variation in abundance across site types, but many lineages displayed fluctuations in abundance across seasons in the RNA dataset, but not the DNA dataset (see Fig. 3). The fungal class Agaricomycetes and the bacterial phyla Actinobacteria, Alphaproteobacteria, Betaproteobacteria, and Crenarchaeota all increased in abundance throughout the season (see Fig. 3). In contrast, the fungal classes Leotiomycetes, Dothideomycetes, and Glomeromycetes and the bacterial phyla Deltaproteobacteria and Acidobacteria decreased in abundance throughout the season (see Fig. 3).

**Figure 2.**
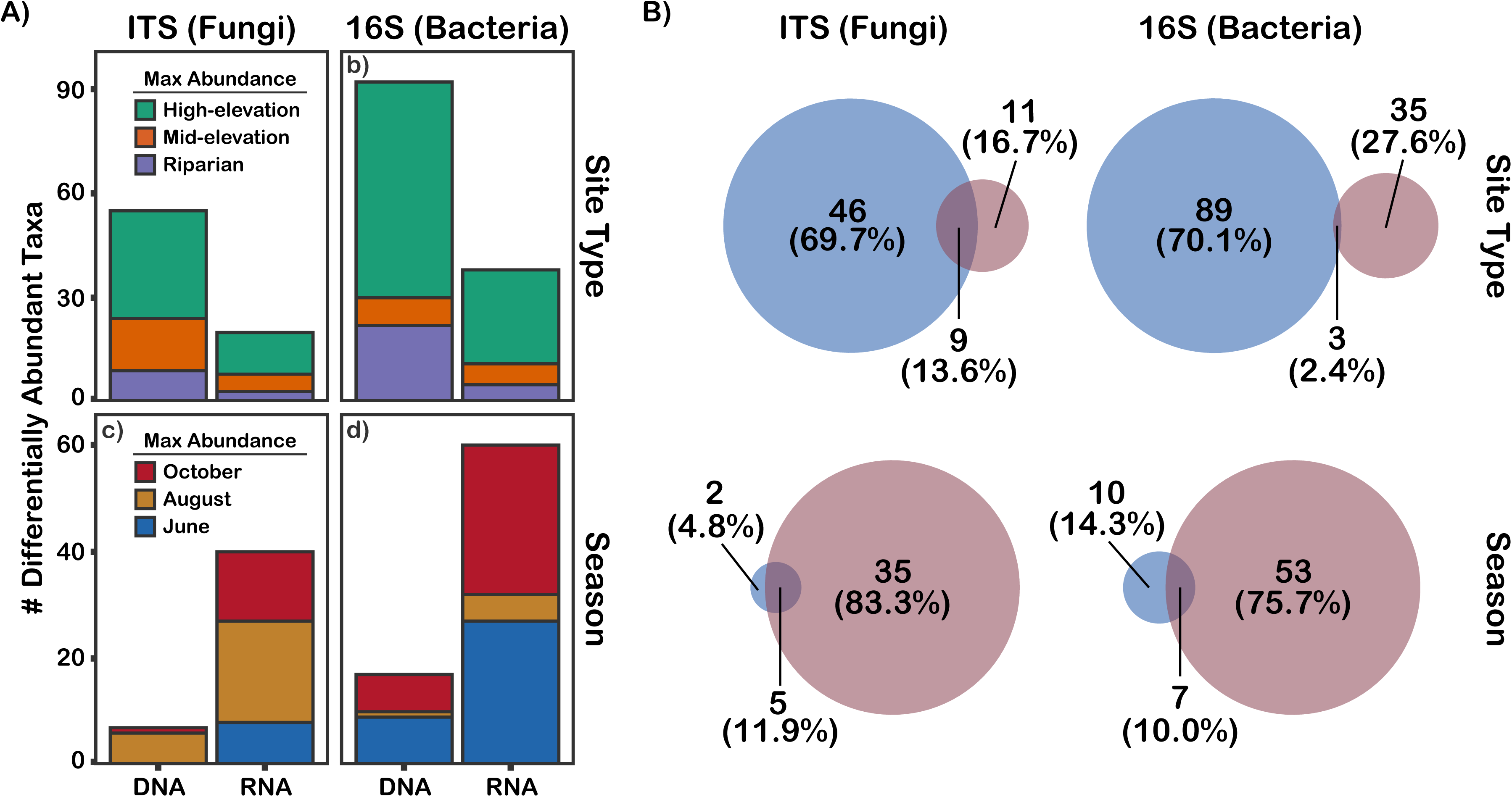
Results of LinDA differential abundance tests for differences in taxon abundance between site types and seasons. A) The number of differentially abundant taxa detected by RNA and DNA metabarcoding and is color-coded by the treatment (site type or season) where the taxa reached their maximum abundance. B) Venn diagrams showing the taxa detected as differentially abundant across site types and seasons by RNA and DNA metabarcoding. The area of the Venn circles is proportional to the number of taxa

**Figure 3.**
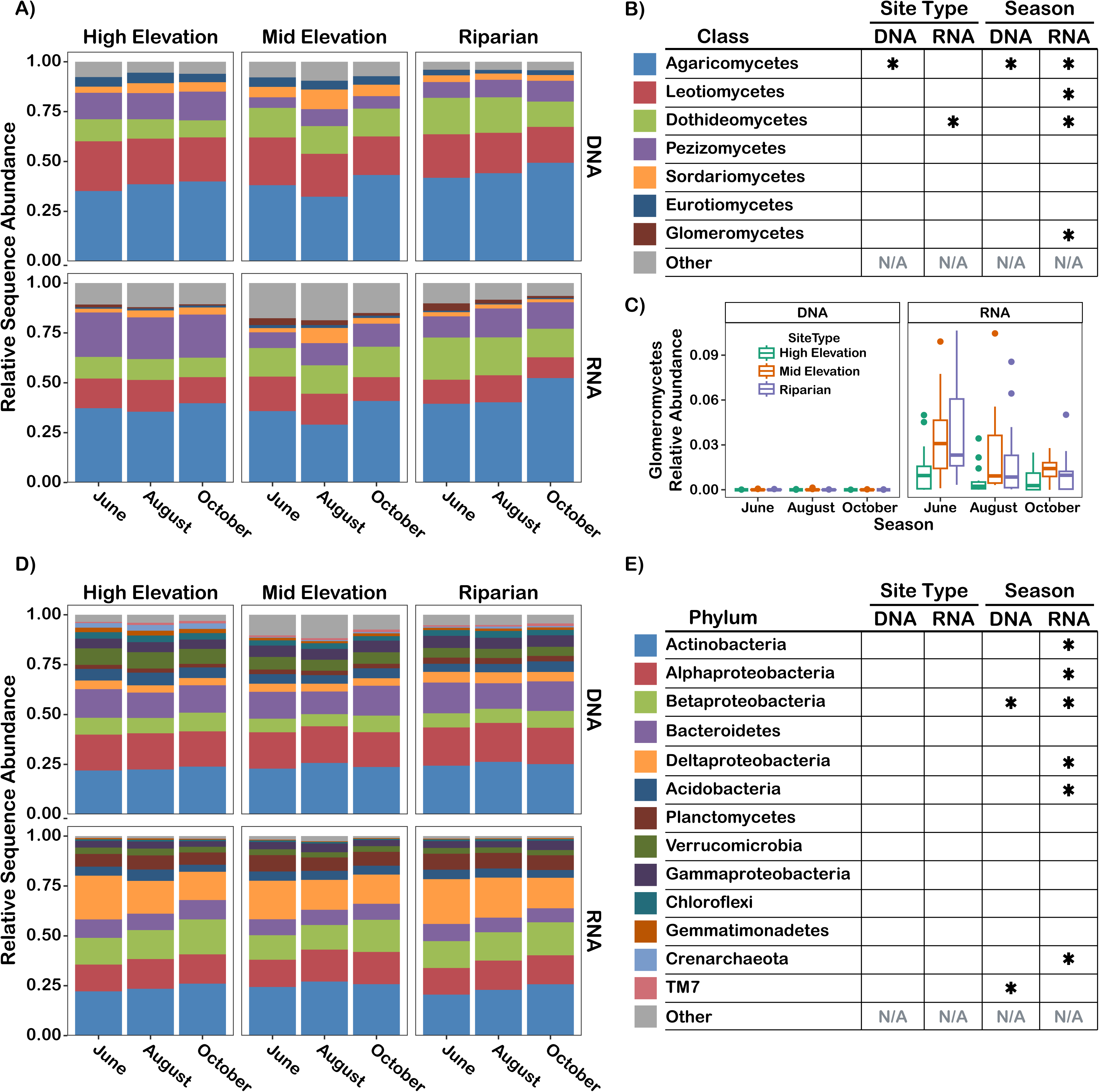
A) Relative sequence abundance of dominant fungal classes (>0.1% relative sequence abundance) across site types and seasons in the DNA and RNA datasets; B) Legend and significance of site type and season effects on the dominant fungal classes; C) Relative abundance of the arbuscular mycorrhizal Glomeromycetes across site types and seasons in the DNA and RNA datasets; D) Relative sequence abundance of dominant bacterial phyla (>0.1% relative sequence abundance) across site types and seasons in the DNA and RNA datasets. Note that for Proteobacteria, data were examined at the class level; E) legend and significance of site type and season effects on the dominant bacterial phyla

Beta diversity analyses (PERMANOVA, beta-dispersion, and W_d_*-tests) indicated that site type had a strong influence on fungal (DNA: R^2^ = 0.11, *P* < 0.001; RNA: R^2^ = 0.07, *P* < 0.001) and bacterial (DNA: R^2^ = 0.11, *P* < 0.001; RNA: R^2^ = 0.07, *P* < 0.001) community composition in the RNA and DNA dataset, while seasonality only had an effect on bacterial RNA community composition (R^2^ = 0.03, *P* = 0.001). Within-site beta dispersion was greatest at riparian sites and lowest at high elevation sites and tended to increase throughout the growing season for both fungal and bacterial communities (see Fig. S5). Additionally, within-site beta dispersion for fungi but not bacteria was greater in the RNA dataset than in the DNA dataset (see Fig. S5; see tables S10, S11), suggesting greater patchiness in the active community than in the total community. In addition to the effects of site type on community composition, we also found significant effects of site and plot on fungal and bacterial composition. Plot effects (R^2^=0.25-0.29) were stronger than both site (R^2^=0.11-0.14) and site type effects (R^2^=0.07-0.11) for DNA and RNA fungal and bacterial datasets, demonstrating that fine scale differences in community composition between microsites within the same stand were maintained across seasons, despite modest seasonal turnover. DNA and RNA metabarcoding both found a general decrease in fungal (*P* < 0.0001; Table S2) and bacterial (*P* = 0.008; Table S3) Shannon diversity throughout the growing season with some site type-specific exceptions (see Fig. 4). Fungal Shannon diversity was the highest (*P* < 0.0001) and most seasonally stable at the high elevation sites, while bacterial Shannon diversity peaked at the Riparian sites (*P* < 0.0001).

**Figure 4.**
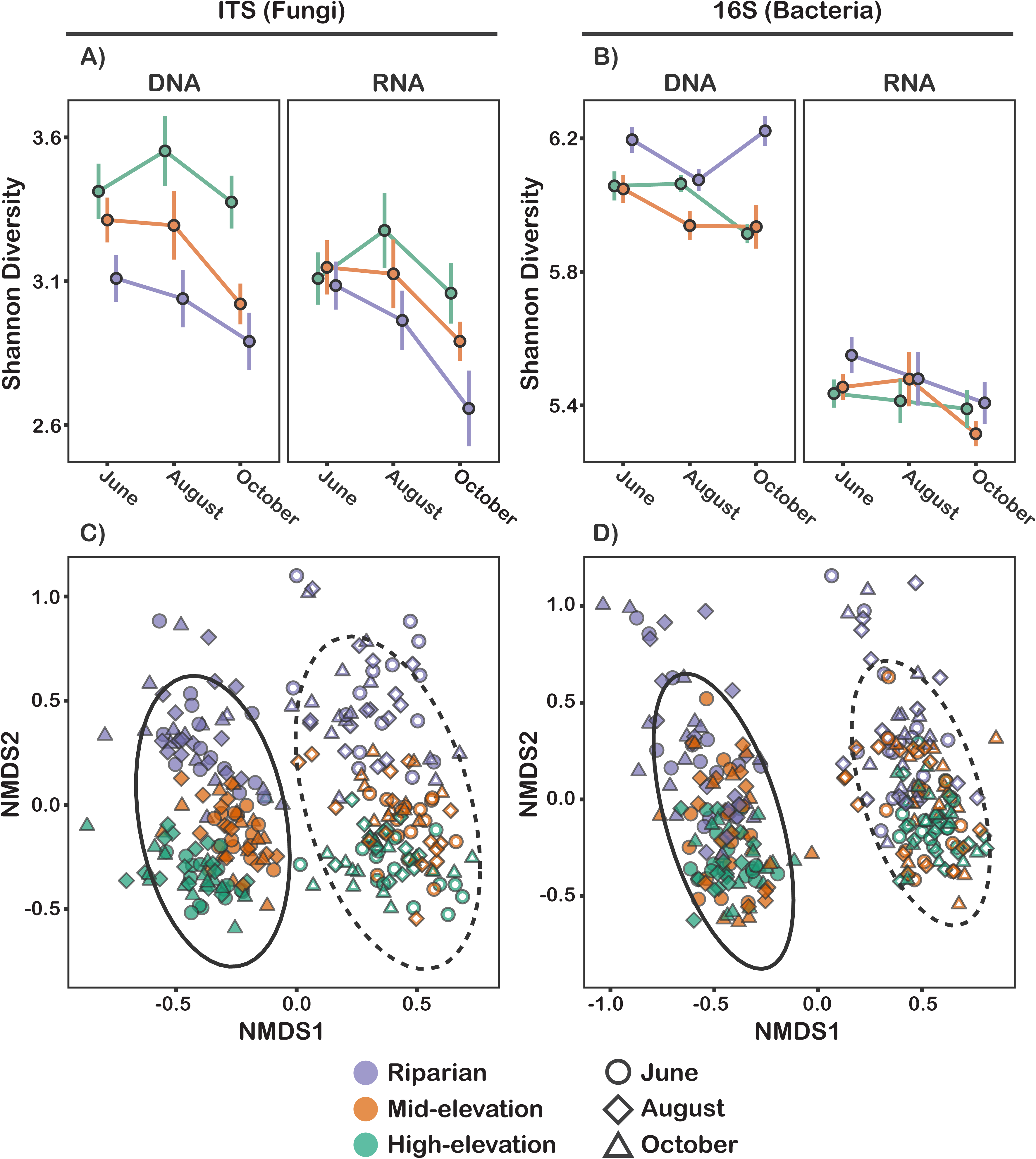
Alpha and beta diversity patterns for 16S and ITS structured by site type, season, and metabarcoding method (RNA vs DNA). a) Fungal communities displayed a general decrease in diversity throughout the season, whereas bacterial communities did not (b). Active communities (detected by RNA) were less diverse than total communities (detected by DNA). c,d) PCoAs of fungal and bacterial community composition show a strong differential between RNA (open points, dashed ellipse) and DNA (close points, solid ellipse) metabarcoding methods. Site type had the greatest effect on community composition, and season had a lesser effect.

We identified environmental factors associated with turnover in community composition, though identification of specific environmental variables responsible for shifts in community composition was made difficult by collinearity between variables. Higher soil pH, base saturation, iron content, and grass cover were consistently associated with riparian community composition (see Fig. S6). In the ITS and 16S DNA datasets, high elevation community composition was associated with a suite of variables associated with cation-rich soils (Ca, Ca, K, Mn, cation exchange capacity, and lime buffer capacity). However, in the RNA datasets, we found weaker predictive power of the environmental variables associated with high-elevation sites, suggesting a decoupling of active community composition from the environment (see Fig. S6).

### Community differences between RNA and DNA datasets

We identified 95 fungal taxa and 538 bacterial taxa that differed in abundance by at least 2x between the DNA and RNA metabarcoding datasets (see Supplemental File S4 for list). This represents 24% of fungal OTUs and 38% of bacterial ASVs that satisfied the filtering requirements of the LinDA tests used for differential abundance testing. Fungal OTUs in the arbuscular mycorrhizal class Glomeromycetes were dramatically underrepresented in the DNA dataset (0.0063%) compared to the RNA dataset (1.84%; P < 0.0001, see Fig. 3C). The bacterial phyla TM7 and Chrenarchaeota were strongly enriched in the DNA dataset, while the Betaproteobacteria and Planctomycetes were strongly enriched in the RNA dataset (see Fig. 3). Community composition was strongly differentiated between the DNA and RNA datasets for both fungi (R^2^ = 0.05, P < 0.001; see table S4) and bacteria (R^2^ = 0.12, P < 0.001; see table S5). Taxon richness was significantly greater in the DNA dataset than in the RNA dataset for fungi (193 vs. 139 OTUs/sample, P < 0.0001; see Table S2) and bacteria (748 vs 453 ASVs/sample, P < 0.0001; see Table S3). On average, 47% (±9%) of fungal OTUs and 34% (±6%) of bacterial ASVs that occurred in a given DNA sample were detected in the corresponding RNA sample suggesting that the active community represented a subset of the total community.

### Identification of an active core microbiome

In addition to identifying microbes within the *Populus* root microbiome that varied by site type and season, we also identified a core *Populus* root microbiome that was present in 95% of samples across site types and seasons. We differentiated between a *total core microbiome* based on the DNA dataset that contained 8 fungal and 51 bacterial taxa and an *active core microbiome* based on the RNA dataset that contained only 2 fungal and 20 bacterial taxa (see Fig. S7). Both total and active fungal core microbiomes were entirely comprised of Ascomycete OTUs. Although six out of the top ten most abundant fungal OTUs were ECM, no ECM fungi met the criteria for either the total or active core microbiome membership because they occurred sporadically across plots. The bacterial active and total core microbiomes were primarily composed of Proteobacteria and Actinobacteria, but also contained some members of the Acidobacteria, Bacteroidetes, Chloroflexi, and Firmicutes phyla. The active core microbiome was generally a subset of the taxa in the total core microbiome (though there were two active core bacterial taxa that were not part of the total core community). Taxa defined as core by DNA metabarcoding were in some cases present at much lower occupancy in the RNA dataset (e.g. the fungus *Exophiala equina* and the bacteria *Mycobacterium* sp., Sinobacteraceae sp., *Phyllobacterium* sp., and EB1017) and in one extreme case (the bacterium *Microlunatus* sp.) could not be detected in the RNA dataset at all.

## Discussion

We hypothesized that 1) active fungal and bacterial communities would be more dynamic through time than total communities, 2) microbial taxa would vary in abundance across ecologically distinct sites and across seasons, and 3) core microbiome members would be highly metabolically active through space and time. We found strong evidence to support hypothesis 1, with a greater number of seasonally differentially abundance microbial taxa detected by RNA metabarcoding than by DNA metabarcoding. We also confirmed hypothesis 2, finding a number of taxa and microbial lineages that varied in abundance through space and time. However, we found mixed support for hypothesis 3, finding that many core taxa were inactive in many of the samples in which they occurred and a small group of core taxa were rarely active.

Our data revealed complex impacts of spatiotemporal environmental variation on the *Populus* microbiome and its activity patterns. Environmental variation associated with site-type (eg. soil chemistry, temperature) was a greater predictor of total microbial community composition than temporal variation associated with seasonal changes (soil temperature, soil moisture, ion fluxes). However, temporal variation in environmental properties associated with seasonal changes had a strong impact on a large number of fungal and bacterial taxa in the active community. There was a large number of fungi and bacteria that exhibited changes in abundance in the active community by season, but not in the total community, suggesting that these changes were due to modulation of metabolic rates rather than fluctuations in population size. Modulation of metabolic activity levels may actually enable the persistence of taxa during unfavorable periods, thus stabilizing microbial communities against environmentally change [8]. We found that the differences in total microbial community composition between site types were possibly driven by soil pH and cation content (Ca, Cd, K, Mn, cation exchange capacity). However, active community composition was also strongly associated with pH, but showed a much weaker association with soil cation content. The finding regarding pH is consistent with a large body of work linking belowground microbial community composition with soil acidity [5, 44], but our findings regarding the decoupling of active community composition from cation content demonstrate potentially distinct environmental controls on total and active microbial community composition.

In addition to a strong ecological imprint of site type on microbial community composition, we found even stronger effects of spatial factors on community composition. Spatial effects were detected at multiple levels, with a significant effect of site and an even stronger effect of plot on both fungal and bacterial community composition. In light of our sampling design with a 2-meter plot size located along transects within stands, this suggests that microsites within forests separated by just tens of meters contain distinct assemblages of microbes that remain distinct despite seasonal changes. Additionally, we found an increase in within-site community dissimilarity for fungi and bacteria throughout the season, suggesting the plots had a tendency to diverge in composition across the growing season. The increases in community dissimilarity (i.e. beta dispersion) and decreases in alpha diversity in the fungal community that we found may be signatures of host stress and dysbiosis due to the loss of a host’s ability to regulate its microbiota and successfully recruit mutualists [45, 46]. Although it is unclear whether *P. tremuloides* trees in our study experienced increased stress throughout the growing season, our data provide tentative evidence for a loss of deterministic assembly later in the growing season, which may be due to a loss of host regulatory effects on the root microbiome or a direct impact of stress on the rhizosphere microbiota.

Although site type was the strongest control on community composition determined at the OTU/ASV level, higher taxonomic level bacterial and fungal lineages responded more strongly in their activity to season than site type. Rhizosphere communities in the early spring displayed a greater activity of the fungal lineages Leotiomycetes and Glomeromycetes (containing arbuscular mycorrhizal fungi) and the bacterial lineage Deltaproteobacteria, but transitioned to increasing activity of the fungal lineage Agaricomycetes and bacterial lineages Actinobacteria and Betaproteobacteria. The Leotiomycetes contain a number of fungal endophytes and the Glomeromycetes are a monophyletic grouping of arbuscular mycorrhizal fungi. Both of these fungal groups are thought to have a lower carbon demand than ectomycorrhizal fungi [47, 48], which may explain their higher activity levels in early spring when trees may be carbon limited due to recent leaf-out [49]. The seasonal changes that we observed in the *Populus* rhizosphere communities may also be driven have been driven by the strong turnover in soil moisture and temperature. Actinobacteria exhibited a modest increase in abundance in October when soil moisture was the lowest and have been previously found to be enriched in plant root microbiomes subjected to experimentally induced drought conditions [50, 51]. Actinobacteria have been hypothesized to have drought tolerance due to the production of quiescent resting spores [17, 52], but our data provides strong evidence that Actinobacteria can maintain metabolic activity during periods of drought which is inconsistent with persistence as dormant spores as a sole mechanism of drought resistance. It is still unclear whether the seasonal changes in microbiome composition and diversity that we observed are indicative of a regular seasonal cycle due to the use of only a single growing season. Another time-series study of *Populus* over multiple growing seasons found a continuous directional change microbiome composition and limited cyclical seasonal patterns, although that study was in an establishing plantation and not a mature stand [53].

Additionally, two of the lineages that varied in activity across seasons (Glomeromycetes and Deltaproteobacteria) were strongly enriched in the active community relative to the total community. Deltaproteobacteria have been previously documented as major components of the *Populus* rhizosphere and root endosphere [29] and have been found to be highly active in uptake of plant carbon during root colonization in rice [54]. Glomeromycetes are frequently overlooked in DNA-based amplicon surveys of plant roots and soils, but our study – along with one previous RNA/DNA metabarcoding study [55] – suggests this plant-mutualistic group is highly metabolically active and temporally dynamic. RNA metabarcoding thus offers a promising alternative to DNA metabarcoding to profile Glomeromycete communities without having to use taxon-specific primers. We did not find evidence that rare taxa are disproportionately active as has been found in a landmark RNA/DNA study of lake ecosystems [9].

In addition to the community members that varied across site types and seasons, we identified a core microbiome that was nearly universally present. Based on DNA metabarcoding, this core consisted of 8 fungal and 51 bacterial taxa. However, we found that many core microbiome members were frequently inactive, with only 2 fungal and 20 bacterial taxa being included in the active core microbiome. Although many ECM fungi were highly abundant, they generally exhibited a patchy occurrence pattern and failed to meet the criteria for core microbiome membership in either the total or active community, which is consistent with another time-series study of *Populus* [26]. The two active core fungal species were the DSE *Hyaloscypha finlandica* and an unidentified Ascomycota sp. that had high sequence similarity to environmental sequences samples from plant roots and rhizosphere soil. Interestingly, we found that a core bacterial ASV identified as a *Mycobacterium* sp. was only active in 17% of samples. Although the *Mycobacterium* strain detected in our study is not closely related to human-derived isolates, another member of the genus that is implicated in Tuberculosis displays a pronounced dormant phase as part of the infection process [56] and the body-inhabiting strain *M. smegmatis* can enter a dormant phase with low rates of RNA synthesis [57]. These findings potentially challenge the understanding of core microbiomes as essential and functionally important members of the microbiome [19]. Instead, we find that they are frequently inactive and might only be classified as core members of the microbiome in DNA metabarcoding studies because of their persistence as dormant propagules or even as relic DNA from dead cells. Various criteria have been considered for core microbiome membership including presence/absence, abundance, and network connectivity [19]. Our study raises the question of whether persistent metabolic activity across a wide array of environmental conditions and through time should also be considered as a core microbiome membership criterion.

To conclude, our study highlights a previously unrecognized layer of temporal dynamism within plant microbiomes, driven by the fluctuating metabolic activity of microbial symbionts. This dynamism likely contributes to the plasticity of genomically-encoded, community-aggregated traits [58], which may offer adaptive advantages to plant hosts in rapidly changing environments and influence ecosystem processes like nutrient cycling and decomposition. This study makes a strong case for time-series molecular studies to focus on active microbial communities to detect community changes. Our work provides a foundation for future research aimed at identifying the environmental stimuli that trigger transitions between microbial dormancy and activity. Additionally, we demonstrate that core microbiota may be often inactive and activity levels might be considered as a key criterion for including taxa in the core microbiome.

## Data Availability

The 16S and ITS reference sequences and raw sequence data generated during the current study have been submitted to GenBank and the Sequence Read Archive under the BioProject PRJNA1224426. Code underlying the analyses and figures presented here is available at https://github.com/jakenash12/PopulusActiveComm

## Acknowledgements

We greatly appreciate the support of the Duke Biology department faculty and staff in conducting this research. We also received tremendous intellectual support and advice from the scientists in the Plant Microbe Interfaces Scientific Focus Area at Oak Ridge National Laboratory. We thank the managers of the Uinta-Wasatch National Forest for allowing us to sample on their lands. Figure 1 was generated in BioRender under an Academic Publication License on agreement number RJ27X779XZ by Vilgalys, R. (2025) and is available on the web at https://BioRender.com/v60e972.

## Description of supplemental files

Supplemental File S1. Supplementary methods describing collection of soil and understory vegetation data and molecular and bioinformatics methods

Supplemental File S2. Supplementary figures S1-S7

Supplemental File S3. Supplementary tables S1-S11

Supplemental File S4. Supplementary data resources including PCR cycle information, environmental metadata, OTU/ASV tables with taxonomic information, and differentially abundant taxa

## Notes

https://www.ncbi.nlm.nih.gov/bioproject/PRJNA1224426

### Competing Interest Statement

The authors have declared no competing interest.

