## Supplemental File S1 for "Time-series RNA metabarcoding of the active *Populus tremuloides* root microbiome reveals hidden temporal dynamics and dormant core members"

**Methods**

*Soil and vegetation metadata collection*

Soil samples were taken from the top 15 cm of soil from each plot in May 2021 and were sent to the University of Georgia soil testing lab (Athens, GA) for soil chemistry analysis (see Fig. S2). We conducted understory vegetation surveys at each plot to measure the percent cover of grasses, forbs, shrubs, ferns, and bare ground using a radial point intercept method. The circular plots were divided into four quadrants and 12 point intercepts were scored in each quadrant (for 48 total intercepts) by dropping a pin flag randomly and recording the vegetation type of the first plant that it contacted. Ten days prior to each sampling date, four pairs of cation and anion ion exchange resin strips (Plant Root Simulators®, Western Ag, Saskatoon, Saskatchewan, Canada) were installed to provide an integrated measure of nutrient availability in the days leading up to sampling. Resin strips were removed when roots were sampled and analyzed by Western Ag following their standard procedures. Teros 11 volumetric soil moisture and temperature sensors (Meter Group, Pullman, WA, USA) were installed at 15 cm depth in the three middle plots at each site to collect data at 15 minute intervals from late May until the final sampling point in October.

Although we had a largely intact time series of soil temperature and moisture data from the sensors, there were windows when individual sensors went offline due to equipment damage. For these windows of missing data, we interpolated data by developing linear models based on the closest online sensors’ data regressed against the sensor of interests’ data and then predicted the missing values based on these linear models. For both soil temperature and moisture, we calculated the following summary statistics – mean average change, minimum mean daily value, maximum mean daily value, average mean daily value, mean diel fluctuation, and range in mean daily value.

*Metabarcoding of plant roots*

The entirety of each fine root sample, including adhering rhizosphere soil, was cryogenically ground in a mortar and pestle with liquid nitrogen. DNA was extracted from 200 mg of ground roots using the DNeasy Powersoil Pro Kit (Qiagen, Hilden, Germany) and RNA from 50 mg of ground roots using the Spectrum™ Plant Total RNA Kit (Sigma-Aldrich, St Louis, MO, USA). RNA was reverse transcribed into cDNA using the Omniscript RT kit (Qiagen) with the primers LR5 (Vilgalys and Hester 1990), ITS4NGR (Cregger et al. 2018; White et al. 1990), and 806R (Lane et al. 1985). Amplicon libraries of 16S and ITS2 libraries were generated from DNA and cDNA using a two-step PCR protocol (see supplemental file S4). Amplicons were cleaned using the Omega Mag-Bind TotalPure NGS kit (Omega, Norcross, GA, USA) and pooled in equimolar quantities. ITS2 and 16S Libraries were sequenced on two separate runs of an Illumina MiSeq with 300 base pair paired end sequencing with v3 chemistry at Duke University’s Center for Genomic and Computational Biology.

Sequence data were processed with QIIME 2 v.2021.11 (Bolyen et al. 2019). 16S forward reads were trimmed to 220 bp and reverse reads were trimmed to 120 bp during amplicon sequence variant (ASV) calling with DADA2 (Callahan et al. 2016). ITS2 paired end reads were merged with PEAR (Zhang et al. 2014) without any trimming prior to processing with DADA2, as it was found to decrease the frequency of low quality bases that result in discarding of reads (Callahan et al. 2016). Merged ITS2 sequences were trimmed using ITSxpress (Rivers et al. 2018). Following ASV calling with DADA2, ITS2 sequences were *de novo* clustered into operational taxonomic units (OTUs) at 97% similarity using VSEARCH (Rognes et al. 2016). We chose to use OTUs for fungal data and ASVs for prokaryotes because OTUs have been shown to outperform ASVs for fungi due to polymorphisms between ITS within-species copies that can result in overestimation of alpha diversity (Tedersoo et al. 2022), while ASVs are considered the gold standard for prokaryotic 16S data (Callahan et al. 2017). We refer to OTUs and ASVs collectively as “taxa” in this manuscript. ITS2 sequences and 16S sequences were assigned taxonomy using the *classify-sklearn* command in QIIME 2 (Pedregosa et al. 2011) with the UNITE v.8.3 99% clustered all eukaryotes (Nilsson et al. 2019) and the Greengenes 13_8 99% (DeSantis et al. 2006) databases, respectively. Fungal OTUs were grouped into guilds using FUNGuild {Nguyen, 2016 #242}.
