## Supplemental File S2 for "Time-series RNA metabarcoding of the active *Populus tremuloides* root microbiome reveals hidden temporal dynamics and dormant core members"

**Supplemental Figure S1.** Mean understory vegetation cover at each site determined with a line point intercept method averaged across the five plots at each site.

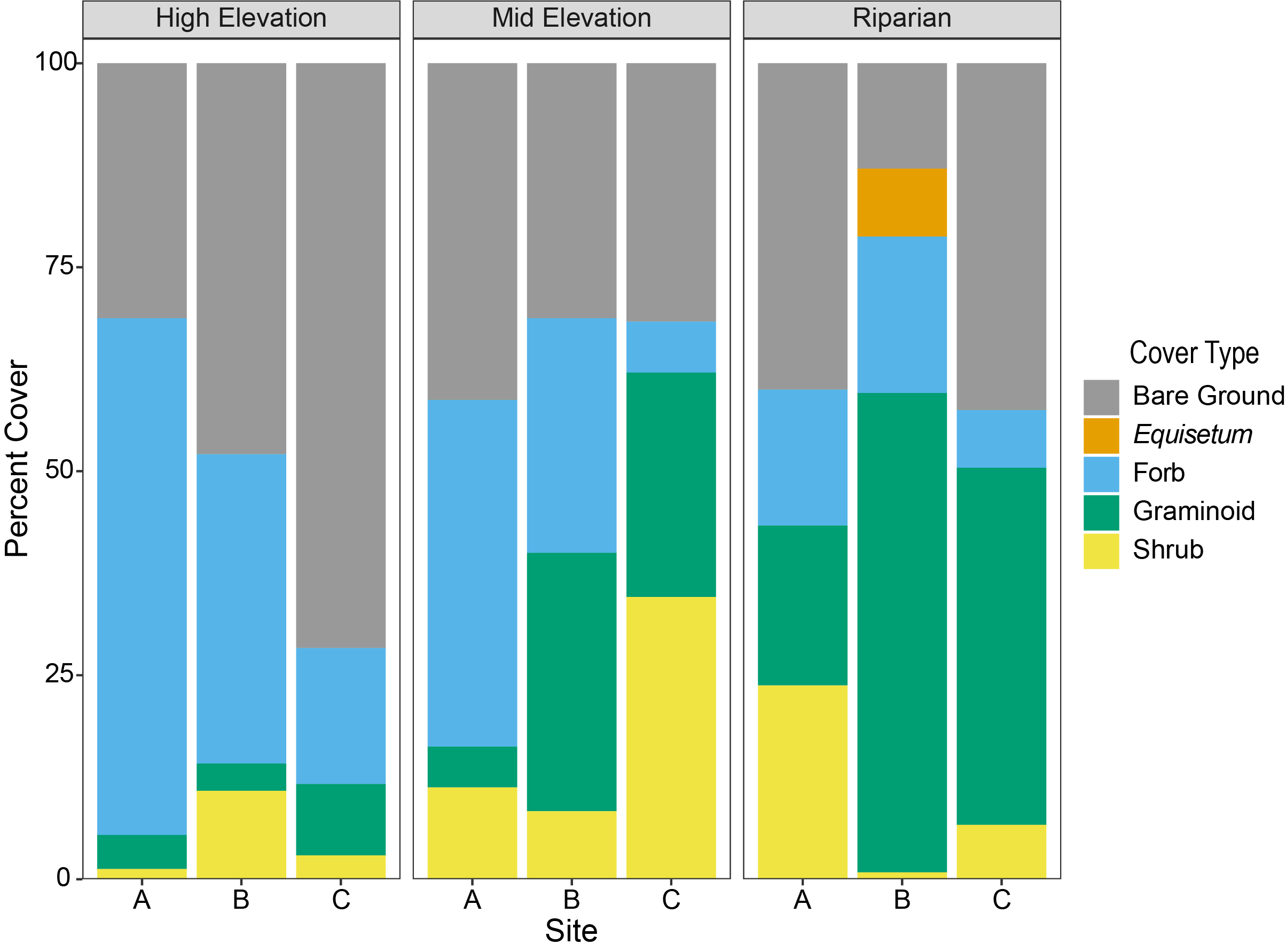

**Supplemental Figure S2.** Soil properties of soils collected at a single timepoint in May 2021. Pairwise significance between site types determined by mixed models is indicated by letters. Plots without letters had no significant pairwise tests.

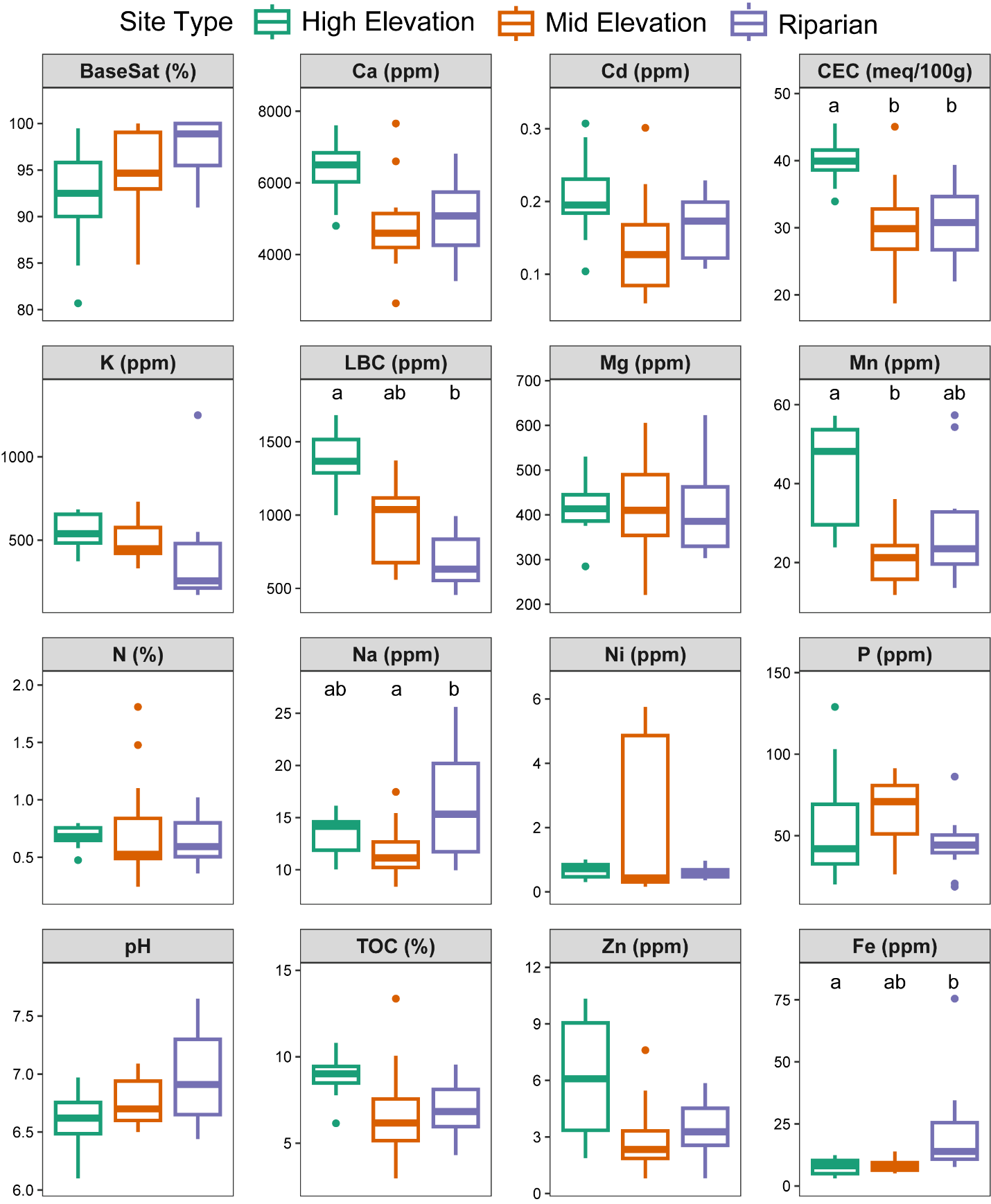

**Supplemental Figure S3.** Average soil moisture and temperature measurements over the course of the growing season (June-October) taken at each plot. Each box represents a different site.

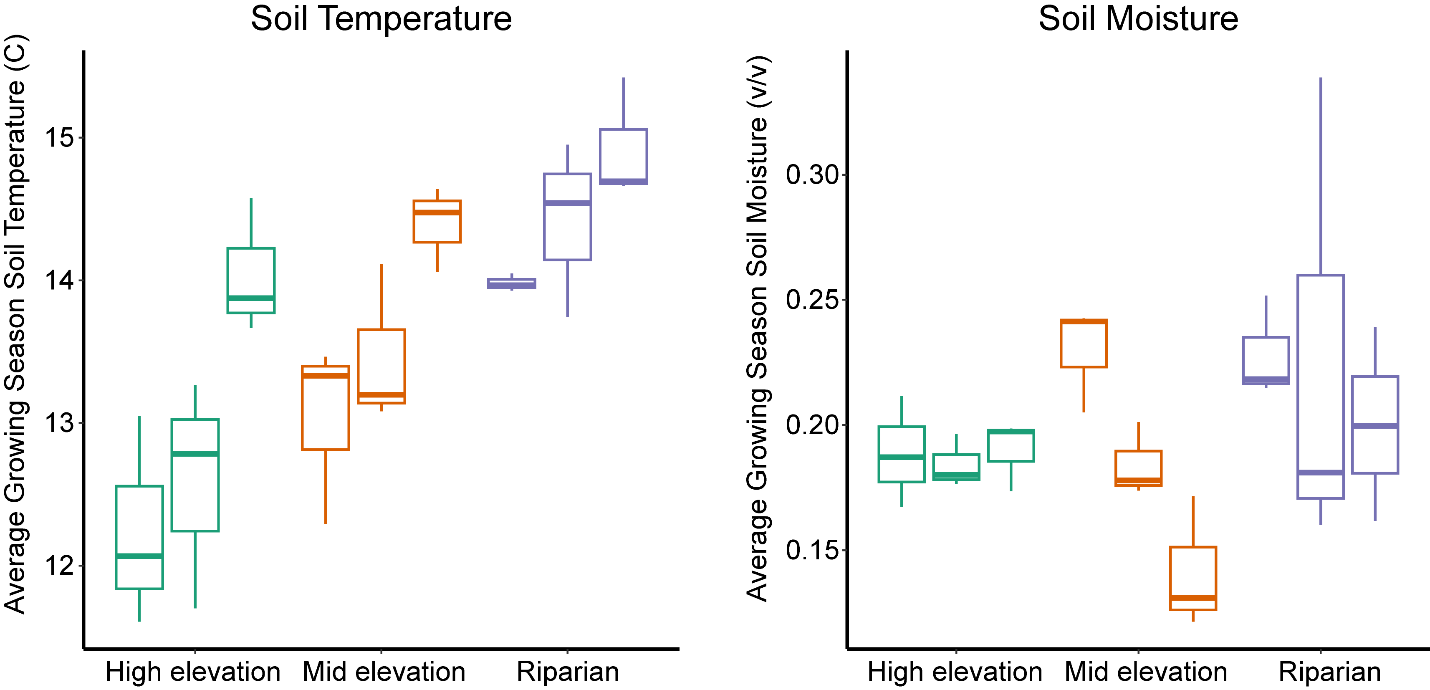

**Supplemental Figure S4.** Ion fluxes determined by plant root simulators across the site types and seasons. Significance of these differences is displayed in Table S1

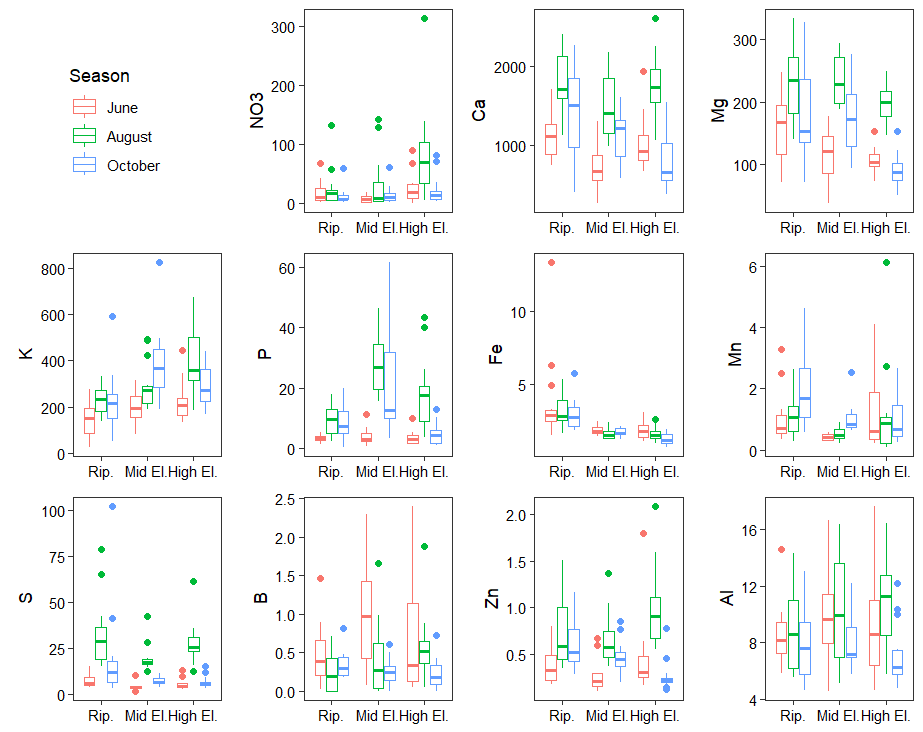

**Supplemental Figure S5.** Mean within-site beta dispersion for the ITS and 16S datasets calculated with Bray-Curtis dissimilarity. Beta dispersion was calculated as the distance to centroid for each unique combination site, season, and barcoding method (DNA vs. RNA). Error bars represent the standard error.

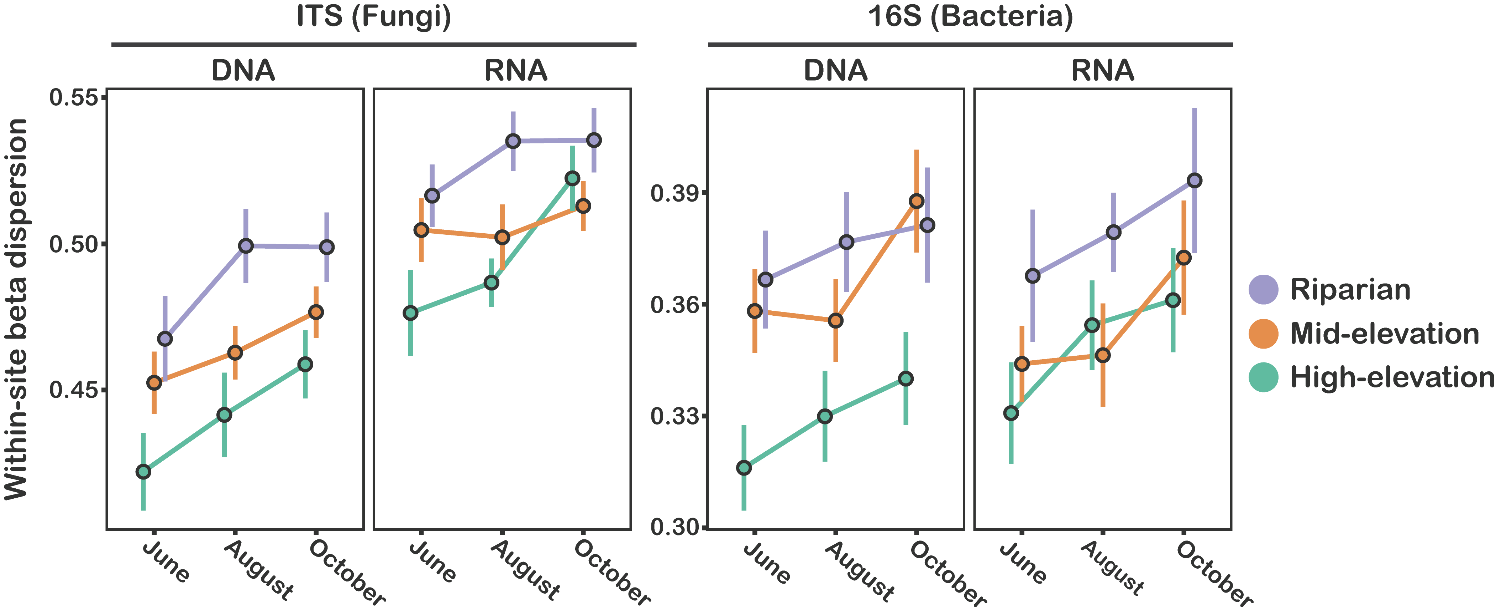

**Supplemental Figure S6.** PCoA biplots with vectors representing environmental variables that were significant predictors of microbial community structure (*P* <0.05) with high explanatory power (R^2^ > 0.3). Separate PCoAs and environmental vector fitting were conducted for the four datasets (ITS DNA, ITS RNA, 16S DNA, 16S RNA). Environmental variables with the suffix “_UGA” are soil chemistry measurements done at the UGA soil testing lab and those with the suffix “_PRS” were measured by soil ion exchange resins that provide and index of ion flux.

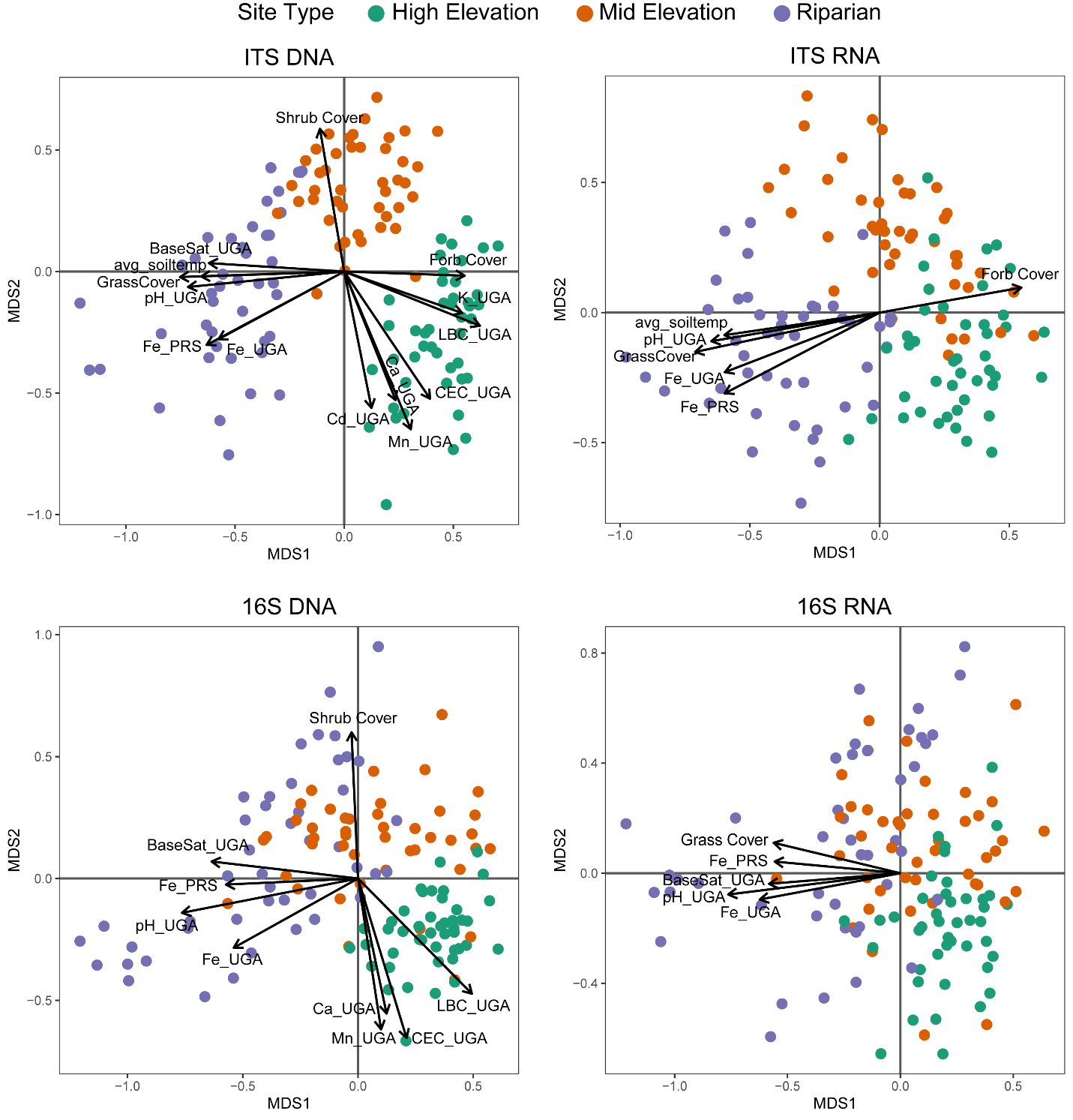

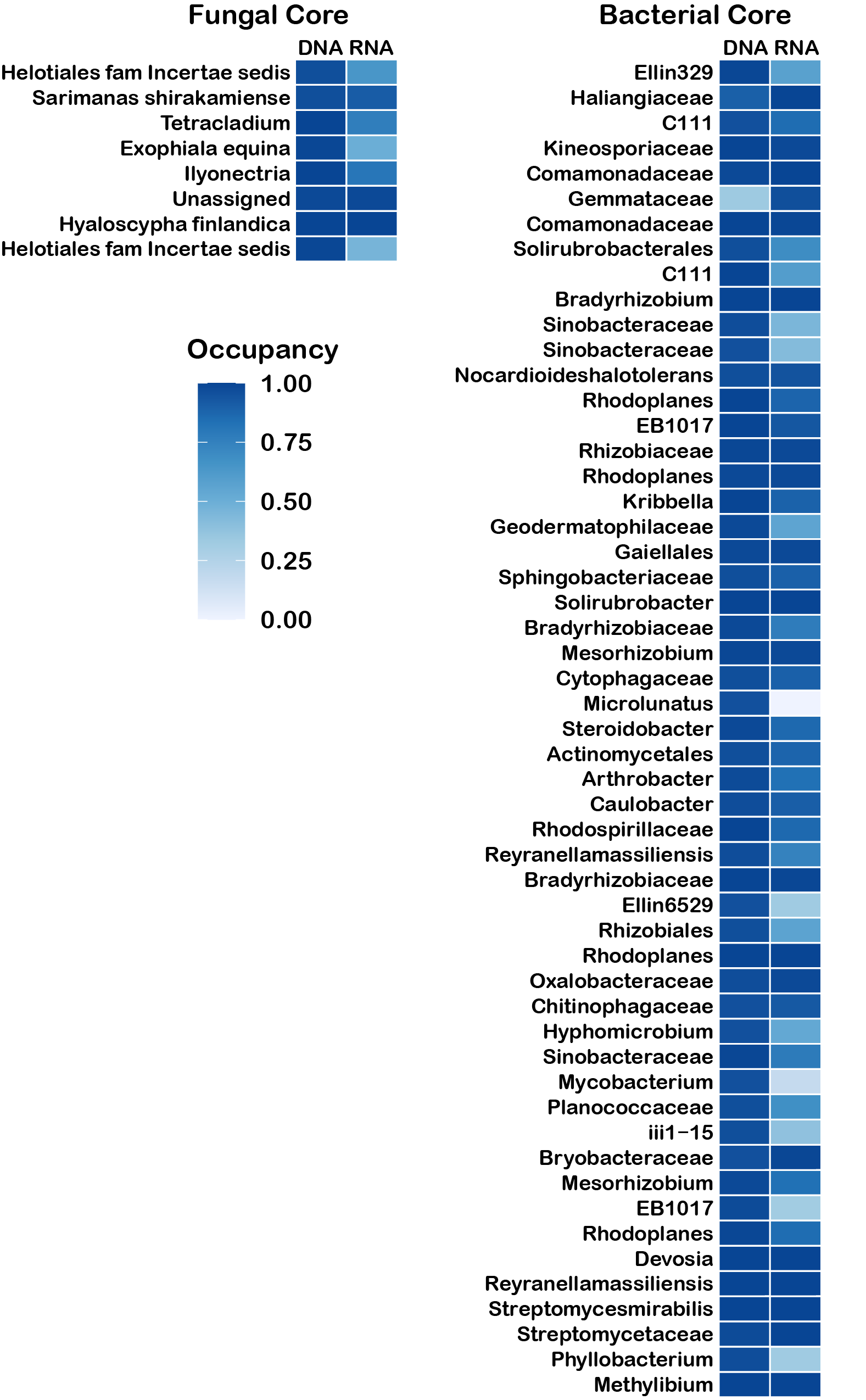
**Supplemental Figure S7.** Occupancy of all fungal and bacterial taxa included in the total and active core communities. Occupancy of 1 indicates presence in all samples, and occupancy of 0 indicates presence in no samples
