## Supplemental File S3 for "Time-series RNA metabarcoding of the active *Populus tremuloides* root microbiome reveals hidden temporal dynamics and dormant core members"

**Supplemental Table S1.** P-values of linear mixed models testing for the effects of site type and season on ion fluxes measured with Plant Root Simulators. Significance is determined by a P < 0.05 and is highlighted in green

| Response | SiteType_p_value | Season_p_value | Interaction_p_value |
| --- | --- | --- | --- |
| NO3 | 0.19403104 | 1.75E-06 | 0.007261109 |
| Ca | 0.389180676 | 3.10E-35 | 2.74E-05 |
| Mg | 0.207338783 | 1.17E-41 | 0.000151054 |
| K | 0.00244165 | 1.94E-08 | 0.002502492 |
| P | 0.036627124 | 4.36E-24 | 6.51E-08 |
| Fe | 1.07E-05 | 0.058754552 | 0.707657484 |
| Mn | 0.457655027 | 0.004357443 | 0.006196071 |
| S | 0.164432686 | 3.20E-21 | 0.076473991 |
| B | 0.07636019 | 1.59E-06 | 0.019869014 |
| Zn | 0.288711842 | 6.49E-14 | 0.000174301 |
| Al | 0.309418868 | 0.000255782 | 0.35814302 |

**Supplemental Table S2**. ANOVA table of linear mixed effects model testing for effects of site type, season, and metabarcoding method (DNA vs RNA) on fungal ITS2 Shannon diversity

| ITS2 (fungal) Shannon Diversity | | | |
| --- | --- | --- | --- |
|  | Chisq | Df | *P* |
| SiteType | 18.5648 | 2 | 9.31E-05 |
| Season | 19.5858 | 2 | 5.59E-05 |
| DNA_RNA | 15.8653 | 1 | 6.80E-05 |
| SiteType:Season | 5.3466 | 4 | 0.2536 |
| SiteType:DNA_RNA | 2.84 | 2 | 0.2417 |
| Season:DNA_RNA | 0.3388 | 2 | 0.8442 |
| SiteType:Season:DNA_RNA | 0.9334 | 4 | 0.9197 |

**Supplemental Table S3**. ANOVA table of linear mixed effects model testing for effects of site type, season, and metabarcoding method (DNA vs RNA) on prokaryotic 16S Shannon diversity

| 16S (prokaryotic) Shannon Diversity | | | |
| --- | --- | --- | --- |
|  | Chisq | Df | *P* |
| SiteType | 18.7065 | 2 | 8.67E-05 |
| Season | 9.7579 | 2 | 0.007605 |
| DNA_RNA | 639.697 | 1 | <2.20E-16 |
| SiteType:Season | 4.1719 | 4 | 0.383248 |
| SiteType:DNA_RNA | 4.8658 | 2 | 0.087782 |
| Season:DNA_RNA | 2.1176 | 2 | 0.346869 |
| SiteType:Season:DNA_RNA | 7.3318 | 4 | 0.119359 |

**Supplemental Table S4.** PERMANOVA table testing for effects of site type, season, and metabarcoding method (DNA vs RNA) on fungal ITS2 community composition using Bray-Curtis dissimilarity with the full dataset. The model uses sequential sum of squares and terms were entered in the order listed below.

| ITS2 (Fungal) PERMANOVA - Full Dataset | | | | | |
| --- | --- | --- | --- | --- | --- |
|  | Df | SumOfSqs | R^2^ | F | *P* |
| SiteType | 2 | 6.782 | 0.06686 | 13.2406 | 0.001 |
| Site | 6 | 9.047 | 0.08918 | 5.8871 | 0.001 |
| Plot | 36 | 22.235 | 0.21918 | 2.4116 | 0.001 |
| Season | 2 | 1.529 | 0.01507 | 2.9854 | 0.001 |
| DNA_RNA | 1 | 5.764 | 0.05682 | 22.5053 | 0.001 |

**Supplemental Table S5.** PERMANOVA table testing for effects of site type, season, and metabarcoding method (DNA vs RNA) on prokaryotic 16S community composition using Bray-Curtis dissimilarity with the full dataset. The model uses sequential sum of squares and terms were entered in the order listed below.

| 16S (Prokaryotic) PERMANOVA - Full Datatset | | | | | |
| --- | --- | --- | --- | --- | --- |
|  | Df | SumOfSqs | R^2^ | F | *P* |
| SiteType | 2 | 5.015 | 0.06085 | 11.7644 | 0.001 |
| Site | 6 | 6.974 | 0.08462 | 5.4528 | 0.001 |
| Plot | 36 | 12.68 | 0.15385 | 1.6524 | 0.001 |
| Season | 2 | 1.045 | 0.01268 | 2.4514 | 0.001 |
| DNA_RNA | 1 | 9.808 | 0.119 | 46.0127 | 0.001 |

**Supplemental Table S6.** PERMANOVA table testing for effects of site type and season on fungal ITS2 community composition using Bray-Curtis dissimilarity with the DNA-only data subset. The model uses sequential sum of squares and terms were entered in the order listed below.

| ITS2 (Fungal) PERMANOVA - DNA Dataset only | | | | | |
| --- | --- | --- | --- | --- | --- |
|  | Df | SumOfSqs | R^2^ | F | *P* |
| SiteType | 2 | 4.724 | 0.10564 | 9.9422 | 0.001 |
| Site | 6 | 5.651 | 0.12637 | 3.9645 | 0.001 |
| Plot | 36 | 13.19 | 0.29494 | 1.5421 | 0.001 |
| Season | 2 | 0.723 | 0.01616 | 1.5205 | 0.001 |

**Supplemental Table S7.** PERMANOVA table testing for effects of site type and season on fungal ITS2 community composition using Bray-Curtis dissimilarity with the RNA-only data subset. The model uses sequential sum of squares and terms were entered in the order listed below.

| ITS2 (Fungal) PERMANOVA - RNA Dataset only | | | | | |
| --- | --- | --- | --- | --- | --- |
|  | Df | SumOfSqs | R^2^ | F | *P* |
| SiteType | 2 | 3.729 | 0.07257 | 6.1412 | 0.001 |
| Site | 6 | 5.411 | 0.10531 | 2.9707 | 0.001 |
| Plot | 36 | 14.5 | 0.2822 | 1.3268 | 0.001 |
| Season | 2 | 1.332 | 0.02592 | 2.194 | 0.001 |

**Supplemental Table S8.** PERMANOVA table testing for effects of site type and season on prokaryotic 16S community composition using Bray-Curtis dissimilarity with the DNA-only data subset. The model uses sequential sum of squares and terms were entered in the order listed below.

| 16S (Prokaryotic) PERMANOVA - DNA Dataset only | | | | | |
| --- | --- | --- | --- | --- | --- |
|  | Df | SumOfSqs | R^2^ | F | *P* |
| SiteType | 2 | 3.865 | 0.10839 | 9.8264 | 0.001 |
| Site | 6 | 4.83 | 0.13543 | 4.0927 | 0.001 |
| Plot | 36 | 9.031 | 0.25323 | 1.2755 | 0.001 |
| Season | 2 | 0.629 | 0.01763 | 1.5985 | 0.003 |

**Supplemental Table S9.** PERMANOVA table testing for effects of site type and season on prokaryotic 16S community composition using Bray-Curtis dissimilarity with the RNA-only data subset. The model uses sequential sum of squares and terms were entered in the order listed below.

| 16S (Prokaryotic) PERMANOVA - RNA Dataset only | | | | | |
| --- | --- | --- | --- | --- | --- |
|  | Df | SumOfSqs | R^2^ | F | *P* |
| SiteType | 2 | 2.748 | 0.07415 | 6.1985 | 0.001 |
| Site | 6 | 4.604 | 0.12422 | 3.4614 | 0.001 |
| Plot | 36 | 9.762 | 0.26342 | 1.2233 | 0.001 |
| Season | 2 | 0.882 | 0.0238 | 1.9898 | 0.001 |

**Supplemental Table S10.** ANOVA table of linear mixed effects model testing for effects of site type, season, and metabarcoding method (DNA vs RNA) on beta dispersion on the ITS2 Bray-Curtis dissimilarity matrix

| ITS2 Beta Dispersion of Bray-Curtis Dissimilarity | | | |
| --- | --- | --- | --- |
|  | Chisq | Df | *P* |
| SiteType | 11.812 | 2 | 0.002723 |
| Time | 31.0868 | 2 | 1.78E-07 |
| DNA_RNA | 131.496 | 1 | < 2.20E-16 |
| SiteType:Time | 7.6189 | 4 | 0.10658 |
| SiteType:DNA_RNA | 2.3646 | 2 | 0.306574 |
| Time:DNA_RNA | 1.9484 | 2 | 0.377496 |
| SiteType:Time:DNA_RNA | 1.4382 | 4 | 0.837528 |

**Supplemental Table S11.** ANOVA table of linear mixed effects model testing for effects of site type, season, and metabarcoding method (DNA vs RNA) on beta dispersion on the 16S Bray-Curtis dissimilarity matrix

| 16S Beta Dispersion of Bray-Curtis Dissimilarity | | | |
| --- | --- | --- | --- |
|  | Chisq | Df | *P* |
| SiteType | 10.3893 | 2 | 0.0055461 |
| Time | 15.0583 | 2 | 5.37E-04 |
| DNA_RNA | 0.5112 | 1 | 4.75E-01 |
| SiteType:Time | 2.4108 | 4 | 0.6606851 |
| SiteType:DNA_RNA | 6.0431 | 2 | 0.0487267 |
| Time:DNA_RNA | 0.2233 | 2 | 0.8943432 |
| SiteType:Time:DNA_RNA | 0.2876 | 4 | 0.9905988 |
